## Supplemental Figures for "Development of the genetic toolkit identifies capsular polysaccharides as a competitive colonization factor in *Ruminococcus gnavus*"

¥¥

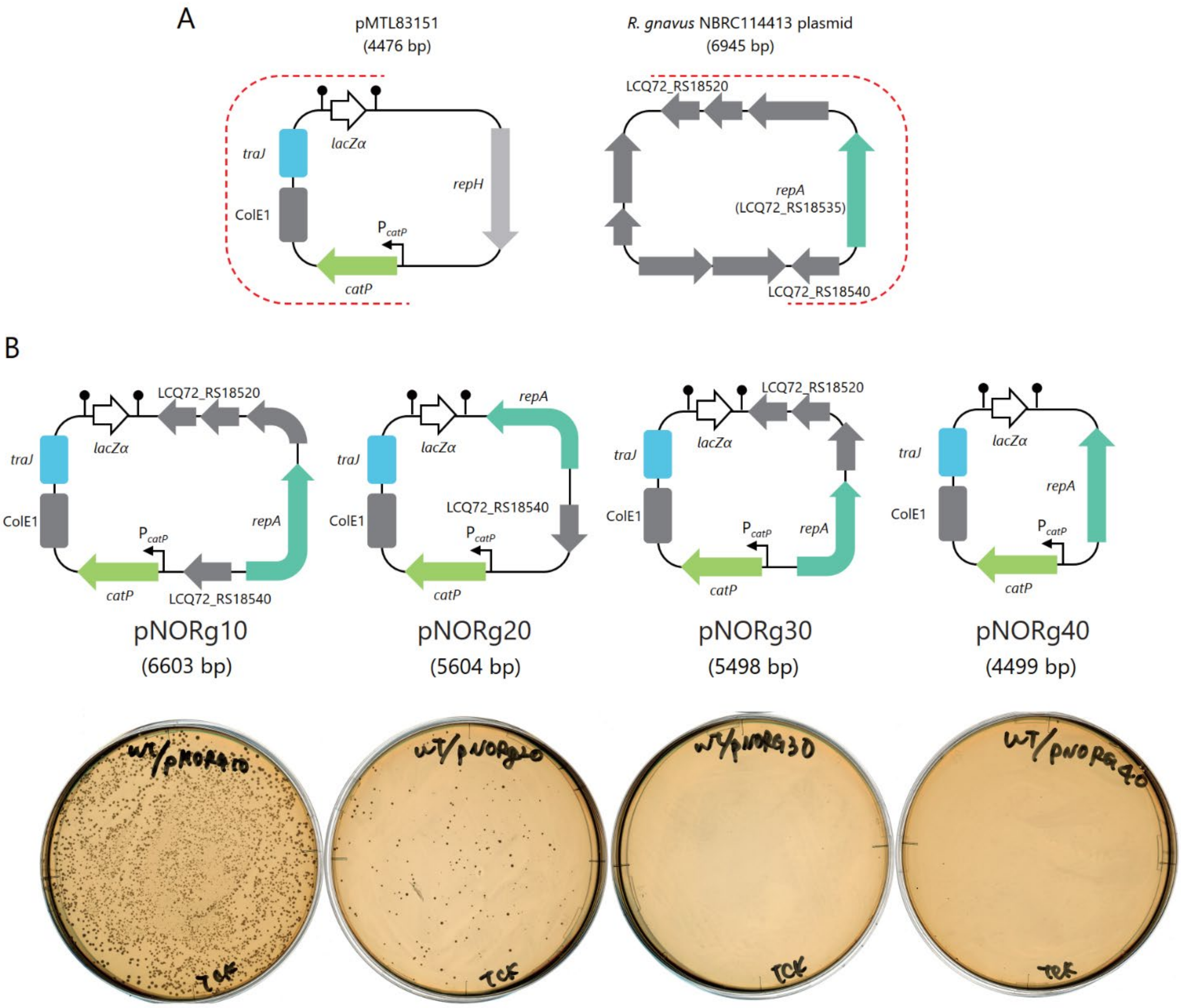


**Supplementary figure 1. Establishment of an *R. gnavus-E. coli* shuttle vector system.**

A. Schematics of pMTL83151 and an indigenous plasmid in *R. gnavus* NBRC 114413. We used the DNA sequence, indicated by red dot lines, for the construction of the shuttle plasmid, pNORg10.

B. Schematics of pNORg plasmid series. The bottom images show colonies that emerged on selective plates after conjugation. pNORg10 represents the highest conjugation efficiency.

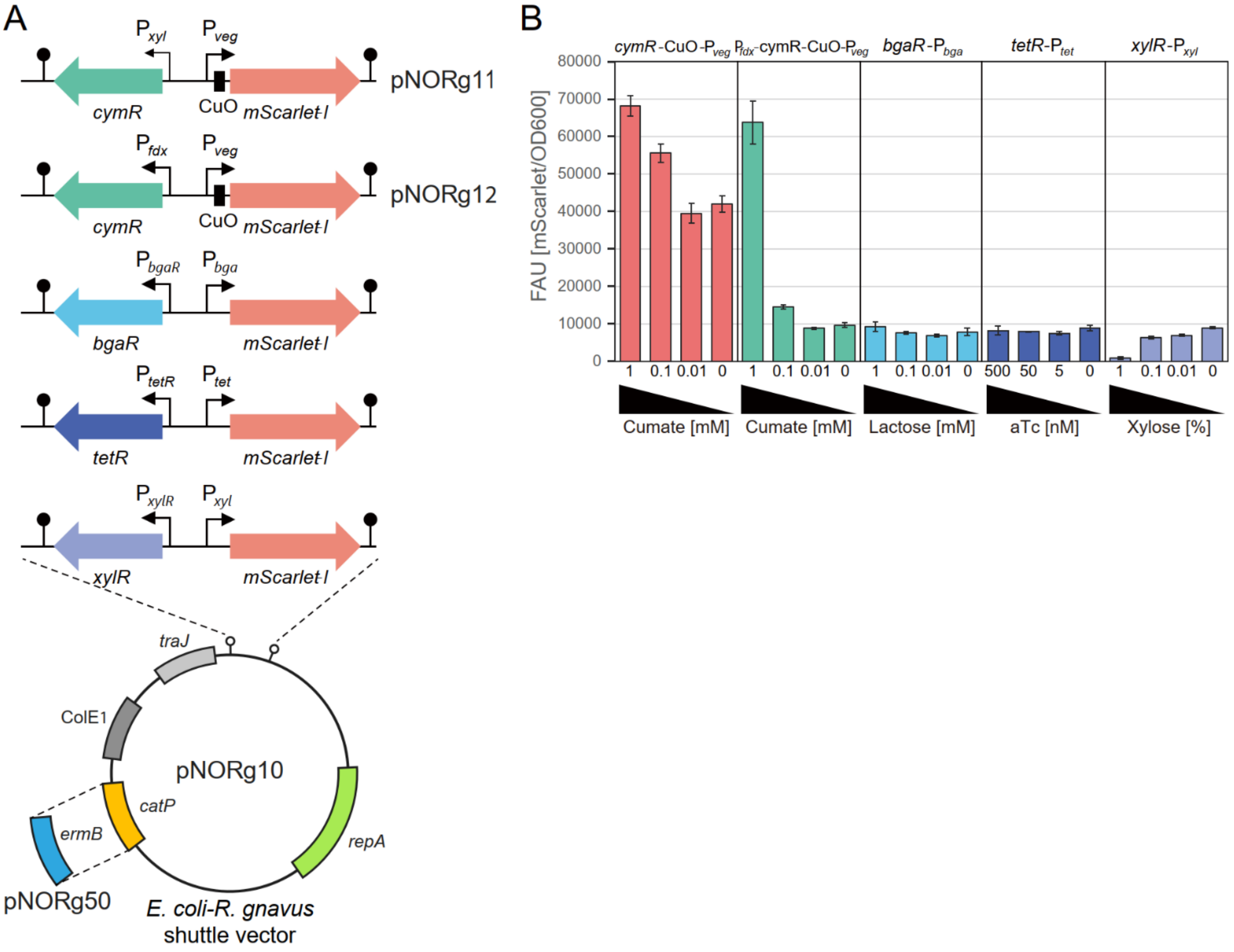


**Supplementary figure 2. Inducible promoter system suitable for *R. gnavus*.**

A. Several inducible promoters were cloned upstream of the *mScarlet-I* gene in pNORg10.

B. Promoter reporter assay using the shuttle plasmids. Cells were grown to the late exponential phase and exposed to oxygen for at least 90 min to facilitate the maturation of the fluorescent protein. The means ± SD of fluorescent intensities normalized by optical density at 600 nm (OD600), obtained from technical triplicates, are shown. The representative results from three independent experiments are indicated.

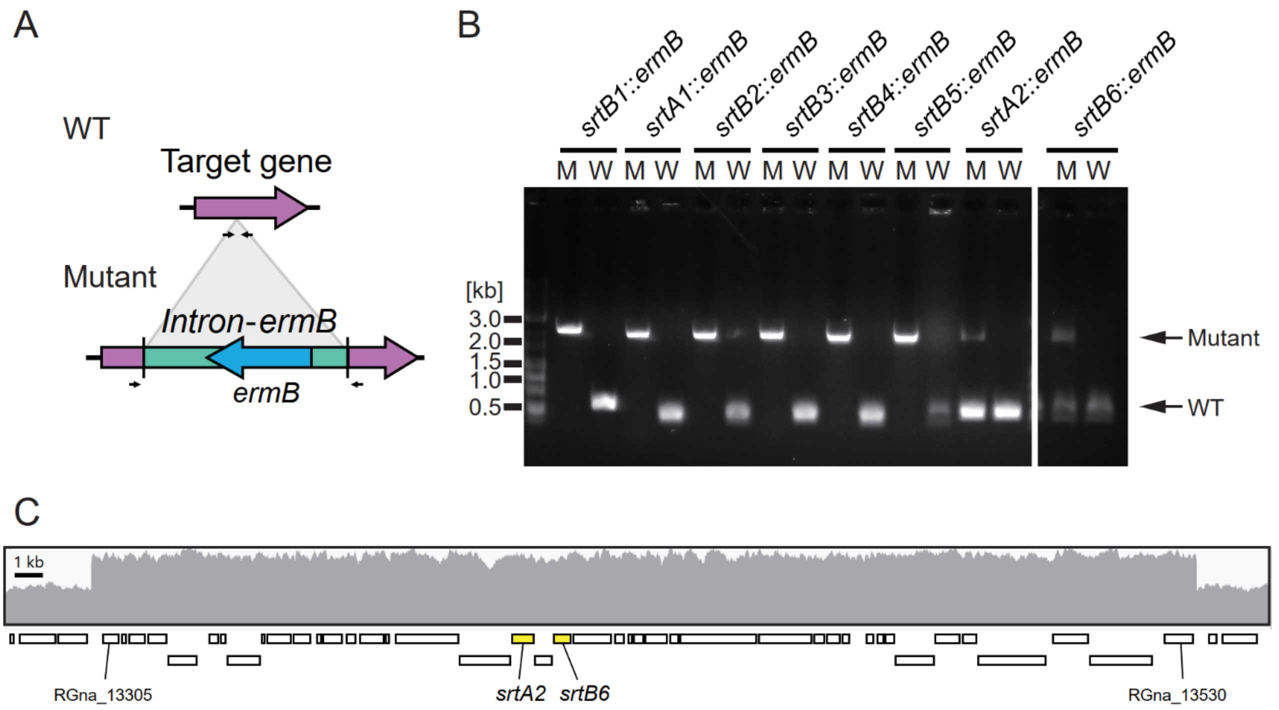


**Supplementary figure 3. Confirmation of sortase gene disruptants in *R. gnavus*.**

A. Schematics of genes targeted by intron. Small arrows indicate the primers used for colony-directed PCR to confirm the gene disruption.

B. Electrophoresis images of the colony-directed PCR. Upper and lower DNA signals indicate PCR products derived from disruptant mutants and the wild-type, respectively.

C. Screenshot of a genome browser. Genomic DNA was isolated from wild-type *R. gnavus* and used for genome resequence analysis. The sequence depth in the genome locus containing RGna_13305 to RGna_13530, which includes the *srtA2* and *srtB6* genes, indicates duplication.

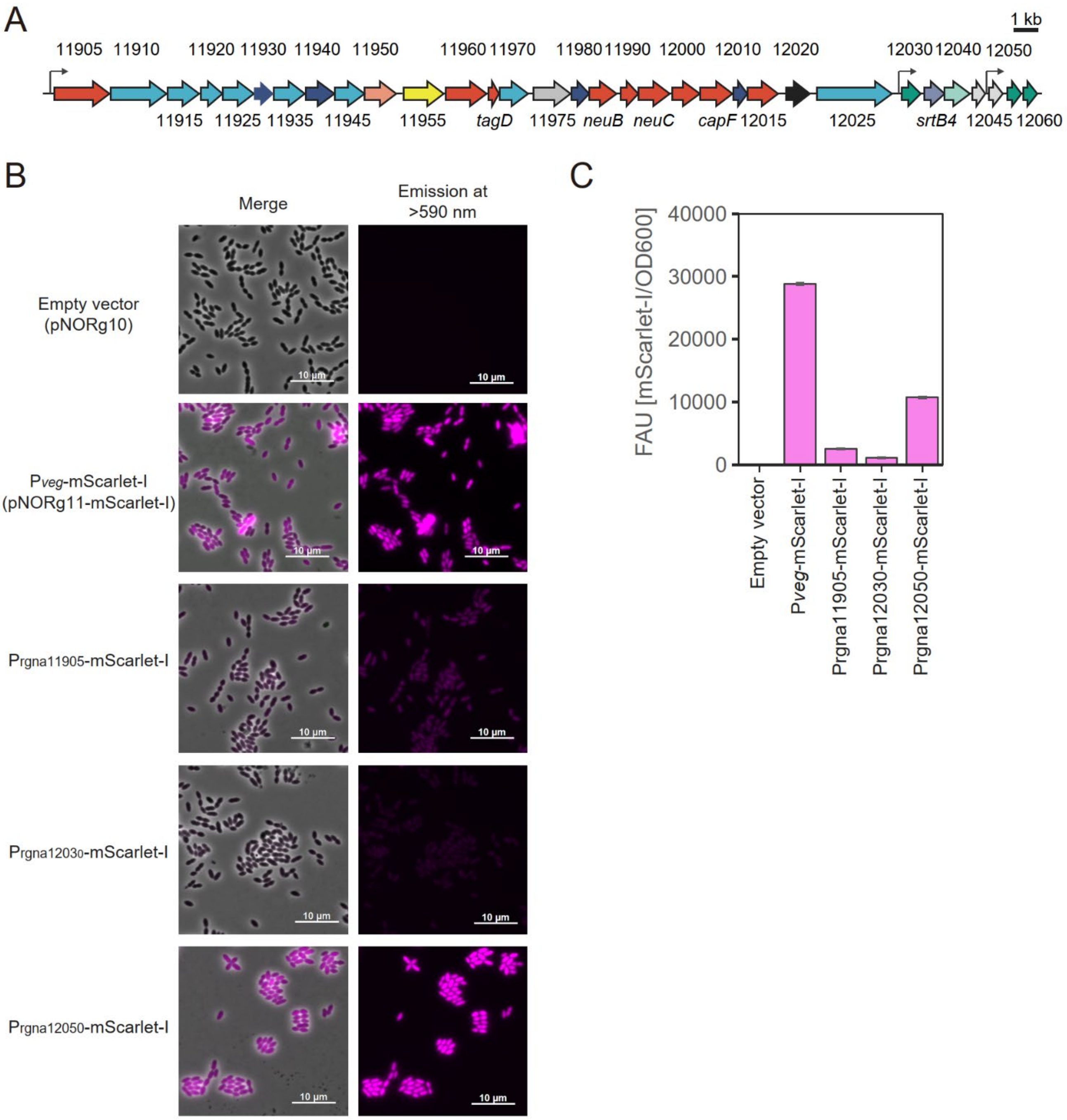


**Supplementary figure 4. Fluorescent reporter analysis of CPS gene promoters in *R. gnavus*.** A. Schematics of the predicted promoters in the CPS gene cluster. Three promoters located upstream of RGna_11905, RGna_12030, and RGna_12050 are shown by bent arrows. B and C. Fluorescent reporter analysis of each promoter. Cells were grown for 6 hours to reach the late-exponential phase and then exposed to oxygen for 90 minutes to mature the fluorescent protein, mScarlet-I. Empty vector and pNORg11-mScarlet-I, including the constitutive promoter $P_{veg}$, were used as negative and positive controls, respectively. Fluorescent microscope images (B) and the means ± SD of fluorescent intensities normalized by optical density at 600 nm (OD600) (C) are shown. The representative results from three independent experiments are indicated.

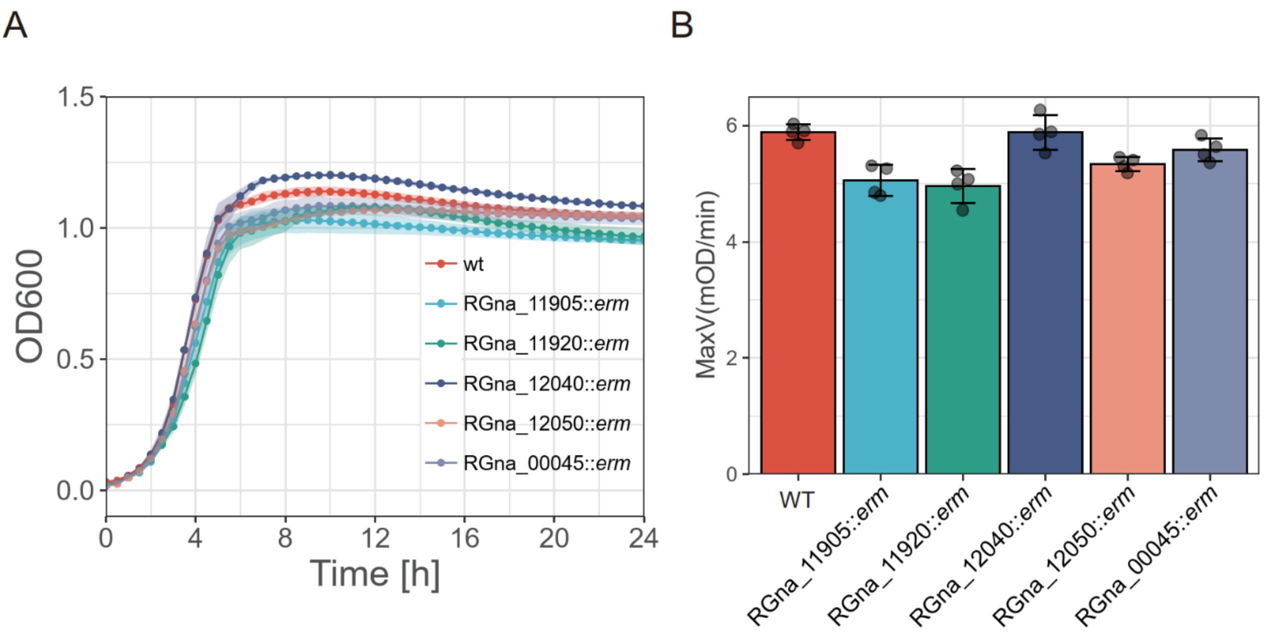


**Supplementary figure 5. Growth curve of CPS gene mutants in *R. gnavus*.**
Growth kinetics (A) and maximum velocities (B) of mutant strains of the CPS gene cluster are shown. The means ± SD obtained from 4 replicates are indicated. We confirmed reproducibility through an independent experiment.

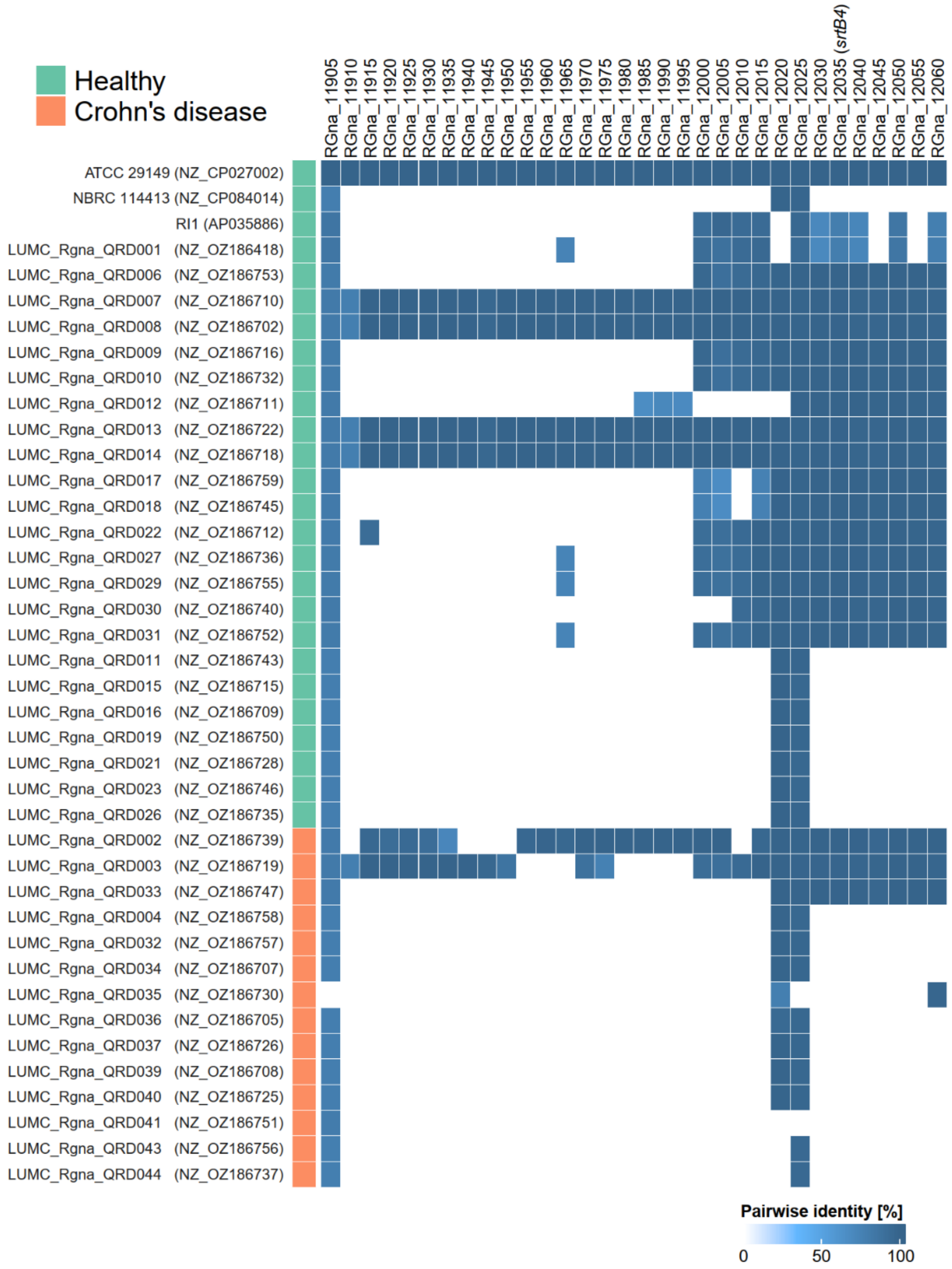


**Supplementary figure 6. Conservation of the entire CPS gene cluster of *R. gnavus* ATCC 29149.**

The conservation and identities of the genes consisting of the entire CPS gene cluster of the ATCC 29149 strain were compared in the genome of isolated strains available in the public database using BLASTn. Forty complete genomes of *R. ganvus* isolates were divided into two groups, highlighted in green and red, representing healthy individuals and patients with Crohn’s disease, respectively. Pairwise identities (%) of each gene are shown as a heatmap.
